# A snapshot of Joshua tree (*Yucca brevifolia* and *Y. jaegeriana*) stand structure in the eastern Mojave Desert of California

**DOI:** 10.64898/2026.01.04.697576

**Authors:** Kathryn A Thomas

**Affiliations:** U.S. Geological Survey, 520 N Park Ave, Suite 221, Tucson, Arizona 85719

**Author notes:** This information product has been peer reviewed and approved for publication as a preprint by the U.S. Geological Survey.

## Abstract

The long-term viability of the iconic Joshua tree of the Mojave Desert is being evaluated. In 2022, we measured the abundance and heights of Joshua tree stems on 62 1000 m^2^ plots in the eastern Mojave Desert of California. The 2022 plots were represented by 33 plots in a population of western Joshua trees (*Yucca brevifolia*) and 29 plots in a population of eastern Joshua trees (*Y. jaegeriana*). Five plots had no Joshua trees; two of which in the western Joshua tree population were destroyed by fire. The 57 plots with Joshua trees supported 627 stems ranging from stems less than 25 cm to mature trees. The plots were examined by abundance and four size classes of the stems, as indicated by their height and indicative of their reproductive status. The western Joshua tree population had more stems (407) than the eastern population (220 stems) although the median difference was not significant. The western population had significantly more stems in pre-reproductive size classes, juvenile (> 25 cm to one-meter, p = 0.001) and sub_adult (> one- to two-meters, p = 0.001), than the eastern population but significantly fewer adult (> two-meters, p < 0.001) stems. The eastern population had significantly greater mean height of adult trees (302 cm +/- 75.3 cm) than the western population (263 cm +/- 52.9 cm) and a larger proportion of stems taller than two-meters (56%) than the western population (39%). In contrast, the western population has 77% of its population in younger size classes (> 25 cm to < 2 meters). These abundance and size class measures alone do not predict whether either population has sufficient natural stand regeneration for long term persistence, but the younger size class structure of the western population suggests greater long-term resilience than for the eastern population.

## Introduction

The persistence of Joshua trees is being evaluated (Wilkening et al. 2020; Shryock et al. 2025), with attention on whether these iconic trees can thrive and regenerate under the variety of threats they currently face. Within the communities in which Josha trees occur, they provide food and shelter to desert wildlife (Borchert and DeFalco 2016, Shryock et al. 2025). The threats that challenge Joshua tree viability are conversion of habitat (urbanization, military training, energy and mineral development); activities that may degrade habitat such as grazing, off-highway vehicles, invasive annual grasses, or altered fire regime; drought and increasing temperatures; and predation and herbivory (refer to Wilkening et al. 2020 review; Smith et al. 2023). The U.S. Fish and Wildlife Service (USFWS; USFWS 2023) has identified the need for additional data to better understand the resiliency of Joshua trees. Such data can help efforts to assess the status of Joshua trees across its range and aid effective decision making for their long-term management and protection (Esque et al. 2015; Wilkening et al. 2020; Shryock et al. 2025).

Joshua trees are perennial long-lived tree-like monocots of the family Asparagaceae, previously Agavaceae. They occur nearly exclusively in the Mojave Desert areas of California, Arizona, Utah, and Nevada (Wilkening et al. 2020). Joshua trees have been considered as one species comprised of two varieties, *Yucca brevifolia* var. *brevifolia* Engelm and *Yucca brevifolia* var. *jaegeriana* McKelvey (Lenz 2007, USFWS 2023). With the recognition that a different species of pollinator is associated with each variety (Pellmyr and Segraves 2003), the varieties were identified as two species of Joshua tree (Lenz 2007), the western Joshua tree (*Y. brevifolia* Engelm.) and the eastern Joshua tree (*Y. jaegeriana (*McKelvey) L.W. Lenz, Fig. 1). Here, we follow the taxonomy recognized by Lenz (2007) and as also accepted by the USFWS (USFWS 2023). The species differ not only in the species of obligate pollinator, but in branching, leaf length, floral and fruiting morphology (Lenz 2007). The adult western Joshua trees are around 6-9 m tall but up to 16 m tall and adult eastern Joshua trees are 3-6 m tall up to 9 m tall (Lenz 2007, Jepson Flora Project 2025.) The distributions of these two species were mapped and described by Esque et al. (2023).

**Fig. 1.**
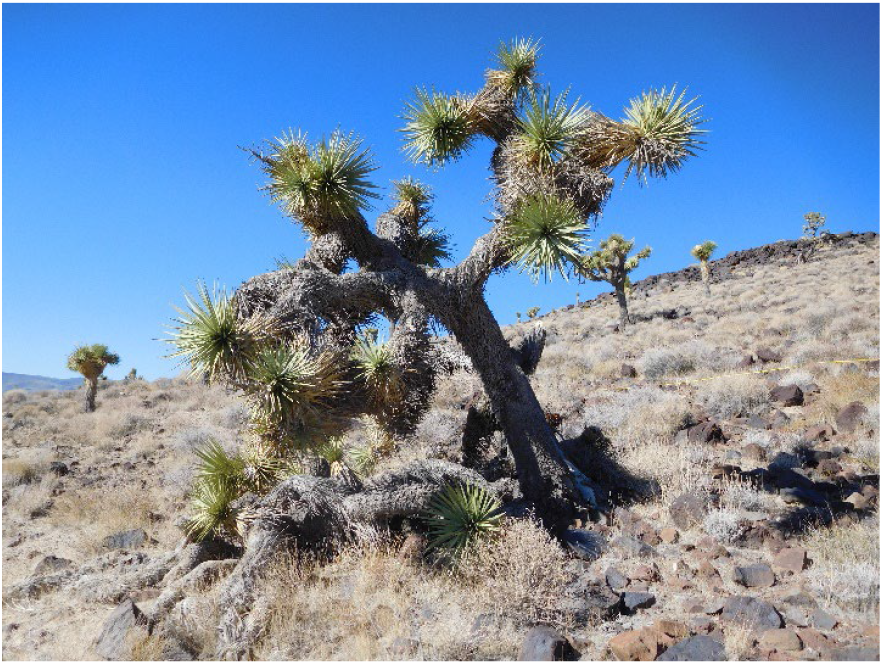
A mature western Joshua tree (*Yucca brevifolia*) in Death Valley National Park. Photo credit: U.S. Geological Survey.

Initiation of the Joshua tree life cycle through sexual reproduction is dependent on flower pollination, seed dispersal, germination and establishment. Success in the seedling stage depends on a complex interaction among optimal precipitation patterns, presence of nurse plants, and minimal herbivory (Reynolds et al. 2012). Joshua trees under 25 cm have been identified as particularly vulnerable (Esque 2015). Stems typically can be reproductive after a height of at least two-meters, but flowering has been observed on plants as short as one-meter. (Esque et al. 2010). Flowering generally happens only periodically, in masting events, associated with increased precipitation years (Yoder et al. 2023). Seed production is the outcome of an obligatory mutualism and requires the presence of the specific pollinating moth, *Tegeticula synthetica* for the western Joshua tree, and *T. antithetica* for the eastern Joshua tree (Smith and Leebens-Mack 2024).

Joshua trees can also regenerate through vegetative growth by clonal sprouts (Shyrock et al. 2014) and or sprouting from fallen trees (D.E. Winkler, U.S. Geological Survey, written comm., Nov 5, 2025). The contribution of clonal sprouting to Joshua tree stand structure and recruitment has not been well studied to date (Todd Esque, U.S. Geological Survey, written comm., Sept. 30, 2025).

Joshua trees are long lived. Multi-year studies of Joshua tree growth rate conducted at various locales in the Mojave Desert have resulted in converging estimates that the tree’s average growth is somewhat greater than 3 cm/yr (Gilliland et al. 2006, 3.75 cm/yr; Esque et al. 2015, 3.12 cm/yr; Cornett 2018, 3.3 cm/yr) but can vary yearly in response to precipitation (Esque et al. 2015) and can exceed this average growth rate in some years (Comanor and Clark 2000). Based on average growth rates and observations of flowering physiology, Esque et al. (2015) estimated a generation time to be 50-70 years at one site in the central Mojave Desert.

In this study we measured the abundance and heights of Joshua trees in plots overlapping or adjacent to plots established in 1997 and 1998. We report here on the resulting measures of Joshua tree stand structure and contrast the measures between the western and eastern populations.

## Methods

In 1997 and 1998, a botanical team made observations on plots in the eastern Mojave Desert of California in support of an extensive vegetation mapping project (Thomas et al. 2004). The approximate plot locations had been pre-selected, using a stratified random sampling framework based on biophysical characteristics. Ultimately, 1105 plots were field located in 1997 and1998, and vegetation and environmental measurements were collected at each. Of the original plots, 174 plots documented the presence of Joshua trees with at least 0.5% cover.

To compensate for the rugged terrain and difficult access to many parts of the eastern Mojave Desert, the 1997 and 1998 sites were allocated in clusters of up to five sites within 1 km of each other. At each of the 1997 and 1998 sites, a plot was established by laying two tapes perpendicular to each other and crossing at the center to form a guide to a circular plot representing an approximately 1000 m^2^ (0.1 ha) circular plot. The 1997 and 1998 plots were not permanently marked, but geographic coordinates were established for the center using a military grade Precision Lightweight GPS Receiver. At each 1997 and 1998 plot the Joshua trees present were documented by cover class and percent cover of the plot, and pictures in each of the four cardinal directions were taken from the plot center. The resulting plot data were reported in Thomas et al. (2018).

Plots for revisit in 2022 were selected from the list of the 1997 and 1998 plots with *Y. brevifolia* documented. The plots selected for 2022 visit were selected from plots on public lands (Bureau of Land Management and national parks) where the field crews could access the plot within an hour as road access was now restricted in some areas open to vehicle access in 1997 and 1998. For each cluster of 1997 and 1998 plots, we selected two plots for revisit or one if two were not available. Given these restrictions, 62 of 174 possible plots were selected for resampling (Fig. 2).

**Fig. 2.**
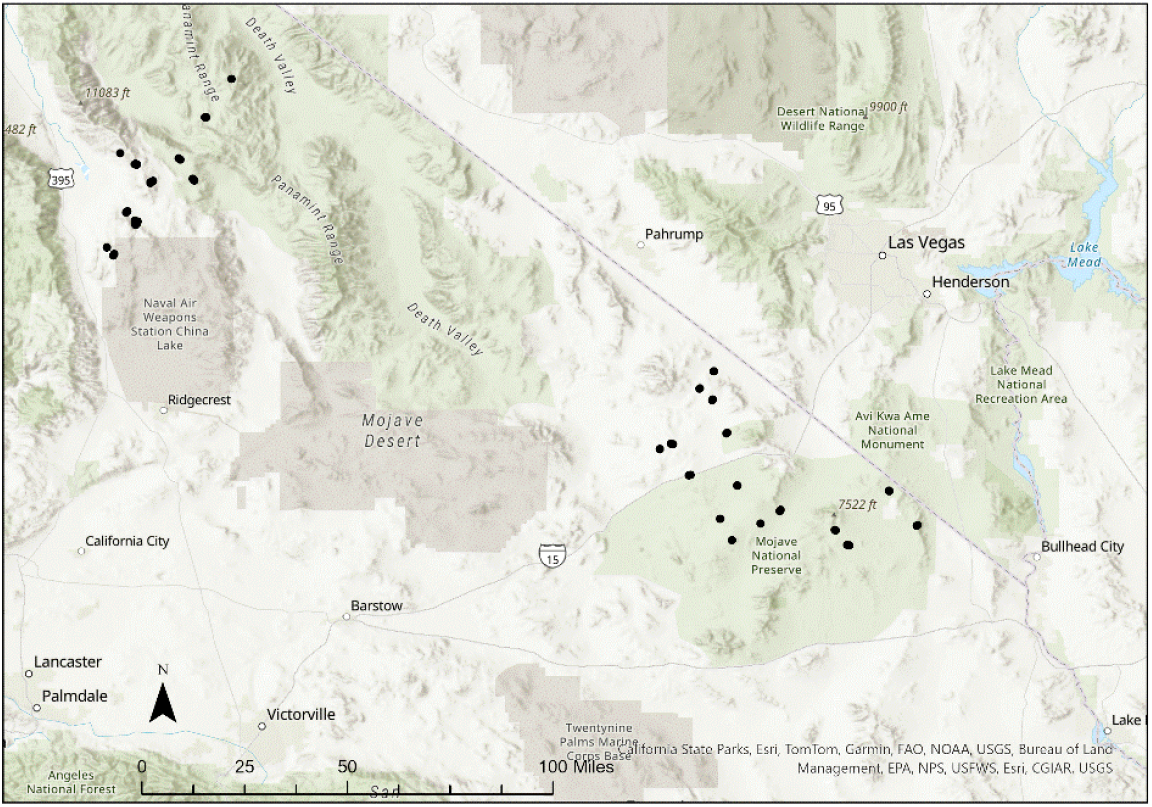
Location of plots (show as black dots) measured in 2022. Sixty-two plots were measured; not all shown uniquely on the map at the scale displayed. The northwest group of plots on the map are western Joshua trees (*Yucca brevifolia*) and the southeast group of plots on the map are eastern Joshua trees (*Yucca jaegeriana*). Base maps are World Topographic Map and World Hillshade provided by ESRI.

Using the same geographic coordinates for each site in the original survey, the field team used contemporary handheld GPS units (Garmin) and copies of photos taken in 1997 and 1998, when available, to relocate the 1997 and 1998 plot locations. Field researchers were instructed to apply the same sampling design used in 1997 and 1998 by laying two tapes at a cross at center so that a 1000 m^2^ circle is delineated by the outer edges of each tape. Flagging was used to identify the outer boundaries of the plot. The exact radius of a 1000 m^2^ plot area was 17.8- and 35.8-meters diameter and was approximated by 18- and 36-meters on the crossed transect tapes

The field team documented all Joshua tree stems within the delineated plot area and measured their height. As small stems were expected to be rare and potentially under shrubs, the field team walked the entire plot and looked under all shrubs for these stems. Photos were taken at each plot. The field team was instructed to stand in the center of the newly established plot and to take pictures in the four cardinal directions, as were the directions given to the 1997 and 1998 field crew.

The field collected data were transferred from paper field sheets to Excel spreadsheets and quality checked by comparing field entered data sheets to the transcribed data to identify transcription mistakes and or data that was not logically consistent. Subsequent comparison of the field photos taken in 2022 with field photos taken in 1997 and 1998 indicated that the plots established in 2022 were offset from the 1997 and 1998 plots, despite the geographical location coordinates being the same. Visual comparison of landscape features in 2022 photos with those in 1997 and 1998 photos indicated that the 2022 plots were in the same vicinity as the 1997 and 1998 plots but did not have the same plot center and little or no overlap with Joshua trees in the 1997 and 1998 photos. As a result, evaluation of survival of Joshua trees measured in 1997 and 1998 was not possible on an individual tree level as originally intended.

Joshua trees measured in 1997 and 1998 were all identified as *Yucca brevifolia* with no variety designation. Using the extensive distribution mapping conducted by Esque et al. (2023), we determined that the northwest group of plots (Fig. 2) were in western Joshua tree population range (*Yucca brevifolia*), and the southeast group were in eastern Joshua tree population range (*Yucca jaegeriana*, Fig. 2). The species only mix and hybridize in a narrow area in Nevada, outside the area of this study (Esque et al. 2023). The USFWS has further designated analysis units within these two population ranges for purposes of assessing the species respective resiliency (USFWS 2023). The plots measured in the western population were in their north analysis unit and the plots measured in the eastern population were in their central analysis unit.

Abundance and height data were collected at the 2022 plots to characterize stand structure within each population (March and October). We measured 62 plots in 2022 (Fig. 2). Fifty-seven of the plots supported 627 Joshua trees. Five plots had no Joshua trees. Two of the plots with no Joshua trees were in the burn area of the 2020 Dome Fire at the Mojave National Preserve that burned 43, 273 acres which had supported large areas of Joshua trees (McAuliffe 2021). The two selected plots had remains of 15 and 11 seemingly dead mature Joshua trees. The other three plots without Joshua trees were in Death Valley National Park (2 plots) and Bureau of Land Management managed land (1 plot). All three had very low Joshua tree cover documented in 1997 and 1998. One of these plots supported two dead stems (one less than 25 cm and the other an adult).

There were 30 measured plots in the western Joshua tree (western) population (excluding the three plots with no live trees) and 27 in the eastern Joshua tree (eastern) population (excluding the two burned plots). All plots were located on public lands managed by the Bureau of Land Management (34 plots: 24 with the western population and 10 with the eastern population), Death Valley National Park (6 plots, western population), and the Mojave National Preserve (17 plots, excluding the burned plots, in the eastern population). The eastern population plots ranged from 1058 to 1595 meters and the western from 899 to 2274 meters.

For analysis of abundance, stems were categorized into four size classes, using height to determine class thresholds, to represent the maturity and vulnerability of the stems based on known vulnerabilities (below 25 cm), estimated growth rates, and estimated heights for sexual maturity (Table 1). A Poisson generalized linear model (GLM) was fit to the plot stem count data for each population to test for dispersion of the count. As the data were highly dispersed, a Wilcoxon rank sum test (W) with continuity correction was used to test abundance between the two populations. The proportions of each population represented in each by size class were tested with Pearson’s chi-squared test (χ2) and subsequent pairwise tests were conducted for each size class with Bonferroni correction. Height for the adult class was tested between populations also using the Wilcoxon rank sum test with continuity correction. The relationship of elevation and abundance by population alone or interacting with population by size class was tested with a negative binomial GLM. All statistical analysis was conducted in R version 4.4.0 (2024). Plots without Joshua trees were not included in these analyses.

**Table 1.** Stem height categories used for Joshua tree size classes. Classes approximate age ranges using growth by year estimates of 3.12 cm/yr (Esque et al. 2015) for low end of range and 3.75 (Gilliland et al. 2006) for the upper end.

| Size Class | Stem Height | Potential age |
| --- | --- | --- |
| Youngest | < 25cm | up to 6.7 to 8 years |
| Juvenile | 25 cm - 1 m | up to 26.7 to 32 years |
| Sub Adult | >1 m to 2 m | up to 53.3 to 64.1 years |
| Adult | >2 m | over 53.3 to 64.1 years |

## Results

The western population supported 407 trees among 30 plots, and the eastern population supported 220 among 27 plots (Table 2). The western population supported 1 to 91 stems of all size classes per plot (mean 13.6 +/-19.9) while the eastern population had plots supporting 1 to 20 stems per plot (mean 8.15 +/-5.6, Table 2). The population median abundances were not significantly different using the Wilcoxon rank sum test (W=422.5, p=0.79).\

**Table 2.** Abundance of Joshua trees is shown organized by population and size class. For comparison, each size class (except All) the mean and standard deviation (Std. Dev.) are for the plots supporting that size class. Proportion represents the proportion of all stems in the entire population represented by that size class. Size classes were defined by height (Table 1). Data for the table comes from Thomas and Winkler (2025).

| Population | Height Category | Num. Plots Containing | Num. Stems | Mean | Std. Dev. | Proportion |
| --- | --- | --- | --- | --- | --- | --- |
| Western | All | 30 | 407 | 13.6 | 19.9 |  |
|  | Youngest | 8 | 22 | 2.8 | 1.8 | 0.05 |
|  | Juvenile | 21 | 126 | 6 | 7.1 | 0.31 |
|  | Sub Adult | 20 | 148 | 7.4 | 11.1 | 0.36 |
|  | Adult | 25 | 111 | 4.4 | 4.9 | 0.27 |
| Eastern | All | 27 | 220 | 8.15 | 5.6 |  |
|  | Youngest | 8 | 10 | 1.3 | 0.5 | 0.05 |
|  | Juvenile | 19 | 38 | 2 | 2 | 0.17 |
|  | Sub_Adult | 21 | 48 | 2.3 | 1.9 | 0.22 |
|  | Adult | 23 | 124 | 4.8 | 3.6 | 0.56 |

Tree abundance between the populations differed significantly by size class (χ2(df =3) =52.343, p<0.001). Pairwise proportion tests by size class showed the difference in abundance for the youngest size class was not significant but was for the juvenile size class (p=0.001), sub_adult (p=0.001), and adult (<0.001) size classes. This difference in size class composition between the populations is illustrated in Fig. 3 where the proportion each size class contributes to the entire population is contrasted between the western and eastern populations. While the youngest size class was similar between the two populations, 77% of the western population was greater than 25 cm and less than two-meters (Table 2, Fig. 3) compared to 39% for the eastern population and 56% of the eastern population was in the adult size class compared to 27% of the western population (Table 2, Fig. 3).

**Fig. 3.**
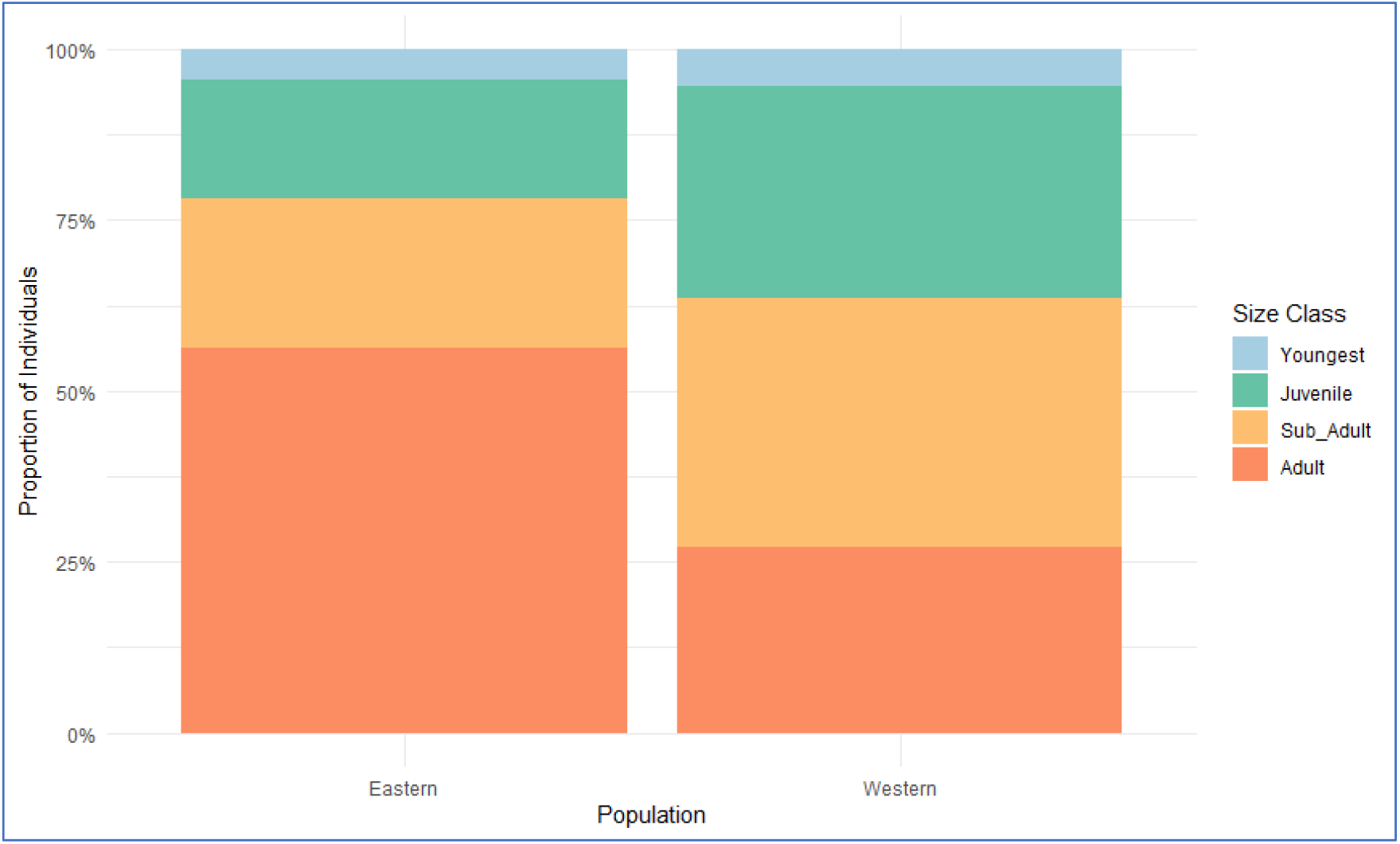
The proportion of each size class of Joshua tree stems within the western and eastern populations. Size classes are defined in Table 2. Data for the figure are available in Thomas and Winkler (2025).

The youngest, juvenile, and sub-adult size classes were bound by height at their lower and upper ends, while adult Joshua trees were only bound at their lower end, two-meters. The mean adult size class height of the eastern population was 302 cm +/-75.3 cm and for the western population 263 +/-52.9 cm. (Fig. 4) and were significantly different (W=9045, p <.001). The tallest tree in the eastern population was 518 cm and in the western population was 439 cm.

Abundance of trees was also tested for their relationship with elevation by population and size class. Elevation alone, by population, or by population and size class was not associated with abundance.

**Fig. 4.**
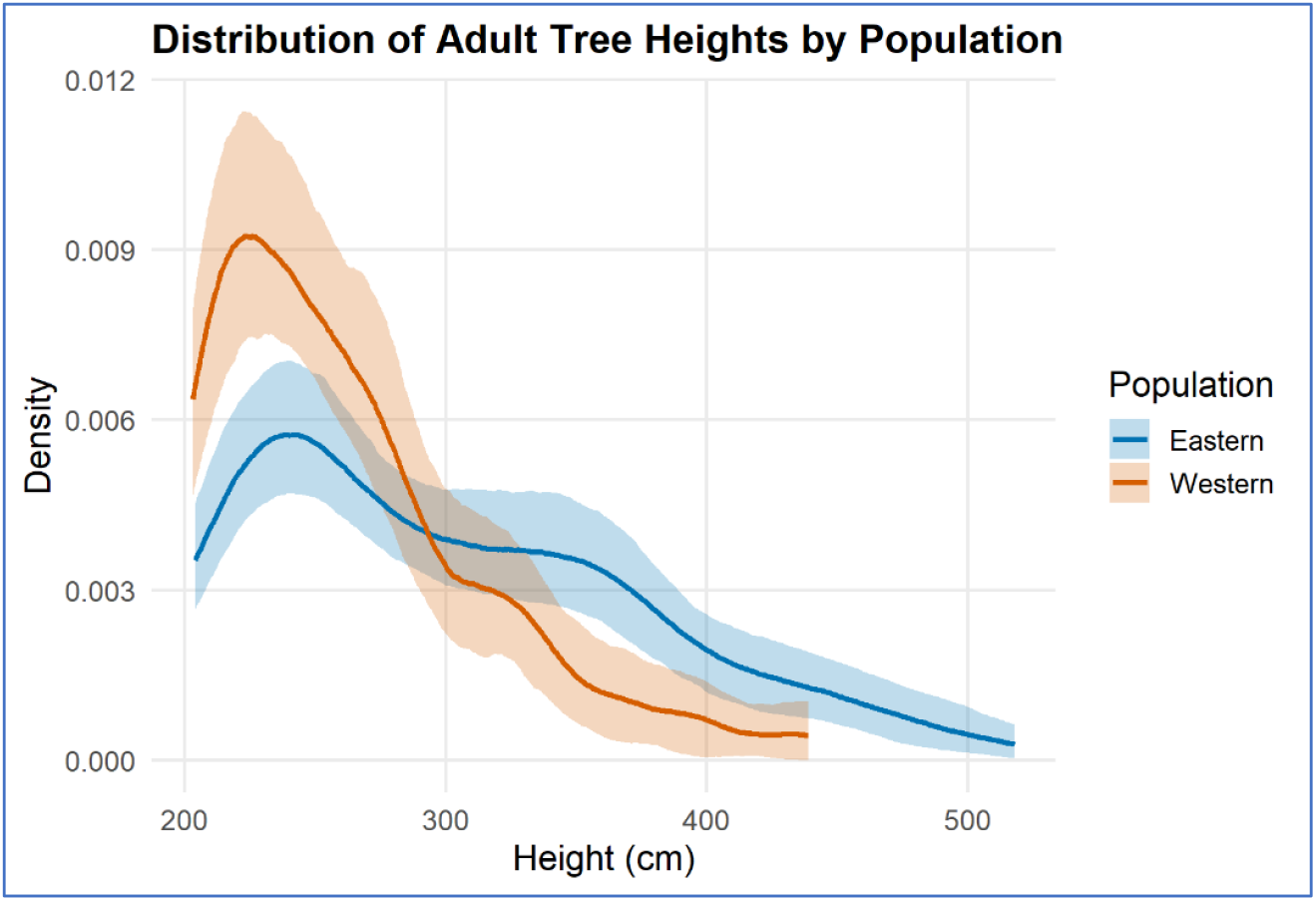
Distribution of adult tree heights for eastern and western populations. The curves represent smoothed height distributions with bootstrapped 95% confidence intervals. The y-axis shows probability density, not counts, with each curve scaled so its total area is one. The image shows the differences in spread and central tendency of adult size class distributions between populations.

## Discussion

This study illustrates differences in Joshua tree stem count (abundance) and structure (size class composition) between a population of western Joshua trees and eastern Joshua trees. The western population had more stems overall and a higher proportion of stems in the 25 cm to two-meter age classes. In contrast, the eastern population reflected an older population with a higher proportion of stems over two-meters in height and with a higher overall average height of the adult size class than the western population.

Stems less than two-meters are generally considered non-reproductive and the two populations differed in two of the three size classes representing these non-reproductive stems. The youngest size class (<= 25 cm) was statistically the same between the populations in numbers of stems and in proportion the class represented among the entire population. In both populations there were only eight plots that supported this size class and with low mean number of stems on each plot. Early establishment for seed produced Joshua tree has been established as a bottleneck (Reynolds et al. 2012) because of the vulnerability of the young stems to environmental pressures. Sexual reproduction is also dependent on the success of pollination of flowering trees and subsequent production of seed. The life history, abundance and distribution dynamics of the two *Tegeticula* moth species that pollinate the different populations is largely unexamined with relation to sexual recruitment dynamics but has been identified as an area where more data would be of benefit to understanding Josha tree recruitment (Wilkening et al. 2020, Smith et al. 2023).

The two other non-reproductive size classes, juvenile (25 cm to one-meter) and sub_adult (> one-to two-meters) were statistically and substantially different between the two populations. The western population had the most stems in these size classes, indicating a higher potential for natural stand regeneration than the eastern population, which had most stems in the adult size class. Survival through sub_adult stage appears to have been favored in the western population compared to the eastern population and could be related to having more favorable precipitation and temperature conditions (Reynolds et al. 2012, Shryock et al. 2014), reduced herbivory pressures (DeFalco and Esque 2010, Esque et al. 2015), or greater opportunities for early growth nurturing by nurse plants (Reynolds et al. 2012). Esque et al. (2023) examined differences in the climate niche of the two species using current distribution of adult trees with climate data from 1980 through 2010 and found that the eastern Joshua tree distribution to be much more influenced than the western population by summer precipitation, deriving from North American monsoon precipitation coming from the east. The eastern population stands measured in this study were at the furthest western edge of *Y. jaegeriana* distribution and potentially recruitment and stem growth and survival could have been lower because of decreases in summer precipitation.

While the western Joshua tree size structure and height ranges imply an overall younger population compared to the eastern population, we do not have systematic metrics on the rate of mortality experienced in either population. It is possible that older trees within the western population have had a higher mortality rate than in the eastern population, but without systematic observations of mortality in each population that possibility is unexplored. In contrast to the mature tree height ranges described by Lenz (2007), the study found the eastern population to be taller on the average than for the western population where Lenz’s species description would have indicated the reverse to be expected.

The field measurements in this study did not differentiate between stems representing young and juvenile size classes originating by seed germination and establishment from stems originating from clonal sprouts. Factors favoring clonal recruitment over seed recruitment are only poorly known and mostly as part of post-fire recruitment (St. Claire et al. 2022) or as proposed as a response to drought stress (Harrower and Gilbert 2021). The size classes used were derived from estimates of reproductive capacity as presented in literature which overall lacked differentiation of the nature of the types of stems measured (clonal or seed produced). However, it has been observed that clonal sprouts, such as post fire, can reproduce in as little as 2 years (USFWS 2023), rather than the up to 32 years early flowering at one-meter height and up to 64 years implied by flowering at two-meter height. No stem height information was provided with this communication, and it is not known if clonal sprouts reach reproductive status earlier than young stems from seeds. Additional data in this study and future studies relating stem height to the clonal or non-clonal nature of the stem could be beneficial to understanding of the natural regeneration capacity of the stands.

We report abundance of Joshua trees in each plot rather than density, although the measurements are comparable at the scale of each plot, 0.1 ha. As we selected plot locations intended to be in the same place as previously measured Joshua trees, but rather were overlapping or adjacent to those locations, extrapolating our measures to one-hectare density counts would have assumed ten times extrapolation of our measurements that was not statistically justified. Regardless recent studies of comparative stand structure have indicated density is highly variable depending on the site and that warmer temperatures are negatively correlated with stand density, potentially affecting the success of early establishment (St. Claire and Hoines 2018). The St. Claire and Hoines (2018) finding implies lower elevations might have lower densities, although lower elevation was not designated by meters in their study. Our analysis of abundance by elevation did not show significant differences by population and age class, although we did have limited samples compared to St. Claire and Hoines (2018).

The original intent of the study was not achieved and the reasons for this can provide guidance for future studies intended to monitor Joshua tree demography through time. For this study, the matching of geographic coordinates was not sufficient for the tree-by-tree matching that would allow assessment of survival of individual trees over the approximate 25-year interval between the repeat visits. The geographic positioning systems used in 1997 and 1998 and 2022 were different, and the lack of match may have been due to unreliable recording of the datum used or other use of the equipment. In retrospect, marked 1997 and 1998 plot centers could have been used for the intended repeat measurements. Alternatively, careful photo matching in the field could have facilitated plot matching, an opportunity largely missed in the 2022 study.

A scientific and management application of stand structure data is estimation of whether a population is stable, increasing or declining. With extremely long-lived plants, and plants that may experience optimal seedling recruitment conditions only at long intervals, gaining sufficient data to make these projections is difficult, a challenge acknowledged by the USFWS (USFWS 2023). Currently there are no population viability models suitable to project if a Joshua tree population is resilient, with sufficient natural stand regeneration for long-term persistence (USFWS 2023). This study’s abundance and size class measures alone do not predict whether either population has sufficient capacity for natural stand regeneration for long term persistence, but the overall younger size class structure of the western population suggests greater long-term resilience than for the eastern population. Elucidation of the long-term status of these populations could benefit from measurements that differentiate clonal from seedling origins of recruitment and with measures of reproductive status that are not size-based estimates but rather based on morphological characteristics of flowering at the tree level. Genetic comparisons between *Y. brevifolia* and *Y. jaegeriana* have been done with respect to their respective pollinators (Smith et al. 2011, Smith et al. 2021) but genomic data on the physiological differences between the species, and in particular their response to environmental conditions have been limited until recently (Heyduk et al. 2025). Additional research could help inform the factors affecting Joshua tree persistence.

## Data availability

Data collected in the field in 2022 are available in Thomas and Winkler (2025).

## Acknowledgments

The author thanks Morgan Andrews, Shannon Lencioni, Natalie Wilson, Daniel Winkler for collection of field data and Daniel Winkler and Angela Hoover for data entry (all with the U.S. Geological Survey). We also thank Tasha La Doux and Jim Andre Sweeney of the Granite Mountains Desert Research Center for field accommodation, assistance with field permitting, and information on road conditions. Similar help with permitting and road conditions were provided by Ambre Chaudoin of Death Valley National Park and Erin McConnell of the Bureau of Land Management. Daniel Winkler, Todd Esque and Emily Palmquist of the U.S. Geological Survey provided useful input on the manuscript. This project was funded by a U.S. Fish and Wildlife Service and U.S. Geological Survey Interagency Agreement 4500154515 and the guidance of Felicia Sirchia. Any use of trade, firm, or product names is for descriptive purposes only and does not imply endorsement by the U.S. Government.

## Notes

https://www.sciencebase.gov/catalog/item/68719d29d4be02732f8798f3

### Competing Interest Statement

The authors have declared no competing interest.

